# Targeted Nano Sized Drug Delivery to Heterogeneous Solid Tumor Microvasculatures: Implications for Immunoliposomes Exhibiting Bystander Killing Effect

**DOI:** 10.1101/2022.10.10.510523

**Authors:** Mohammad Amin Abazari, Madjid Soltani, Farshad Moradi Kashkooli

## Abstract

Targeted drug delivery to cancer cells utilizing antibodies against oncogenic cell-surface receptors is an emerging therapeutical approach. Here, we developed a computational framework to evaluate the treatment efficacy of free Doxorubicin (Dox) and immunoliposome at different stages of vascular solid tumors. Firstly, three stages of vascularized tumors with different microvascular densities (MVDs) are generated using mathematical modeling of tumor-induced angiogenesis. Secondly, the fluid flow in vascular and interstitial spaces is calculated. Ultimately, convection-diffusion-reaction equations governing on classical chemotherapy (stand-alone Dox) and immunochemotherapy (drug-loaded nanoparticles) are separately solved to calculate the spatiotemporal concentrations of different therapeutic agents. The present model considers the key processes in targeted drug delivery, including association/disassociation of payloads to cell receptors, cellular internalization, linker cleavage, intracellular drug release, and bystander-killing effect. Our results show that reducing MVD decreases the interstitial fluid pressure, allowing higher rates of the drug to enter the tumor microenvironment. Also, immunoliposomes exhibiting bystander-killing effect yield higher drug internalization, which supports a higher intracellular Dox concentration during immunochemotherapy. Bystander-killing effect alongside intracellular Dox release and persistence of immunoliposomes within tumor over a longer period lead to more homogeneous drug distribution and a much greater fraction of killed cancer cells than classical chemotherapy. Our findings also demonstrate drug transport at tumor microvascular networks is increased by decreasing MVD, leading to better treatment outcomes. Present results can be used to improve the treatment efficacy of drug delivery at different stages of vascular tumors.

## I. INTRODUCTION

Cancer is known as the second major cause of death worldwide ^1^. Chemotherapy, the most common cancer treatment method, is the use of small-sized high cytotoxic drugs such as Doxorubicin (Dox) to damage rapidly dividing cells, but this treatment results in unwanted side effects on healthy tissues. Novel clinical/preclinical treatment methods alongside computational modeling have recently been developed to enhance the accuracy of cancer diagnosis and the efficacy of drug delivery systems ^1, 2^. Targeted delivery of engineered nanoparticles (NPs) to cancer cells has been widely exploited to decrease the side effect of classical chemotherapy while improving their efficacy simultaneously. It is well-known that NPs are readily capable of accumulating in tumors via the enhanced permeability and retention (EPR) effect, originating from hyperpermeable vasculatures and defective lymphatic systems in tumor tissues ^3, 4^. However, NP delivery via the EPR effect (also known as passive targeting) has shown that only about 0.7% of NPs reaches the targeted tumor site, leading to a heterogeneous distribution of drugs with an accumulation principally at the perivascular regions owing to a poor penetration depth in the tumor microenvironment (TME) ^5, 6^. This limited penetration of NPs might be originated from some tumor physiological barriers preventing effective diffusion of NPs, including elevated interstitial fluid pressure (IFP), dense extracellular matrix (ECM), and heterogeneous microvascular network induced by tumor angiogenesis ^2, 5-8^. Therefore, to achieve higher treatment efficacy, there is an urgent need for novel strategies that can effectively transport anticancer agents deep into the tumor site.

Immunotherapy has now been combined with cytotoxic chemotherapy as a capable treatment strategy for solid tumors, and also cancer active targeting has contributed most significantly to this burgeoning area ^9, 10^. Active targeting with surface-engineered NPs includes the attachment of a targeting ligand (e.g., antibody fragments) to the surface of NPs, which when administered to the ECM tracks and targets the specific receptors over-expressed in target cancer cells ^1, 9^. Nano-liposomes are the most widely applied clinically, due to their high biocompatibility, ease of surface modification, pharmacokinetic profile, and pharmacological characteristics, as well as longer circulation time after surface modification ^1, 10-12^. Antibody-targeted nano-liposomes (further known as immunoliposomes) not only enhance the internalization of liposomes into the cancer cells but also minimize the nonspecific distribution of cytotoxic drugs to undesired tissues ^10-12^. Since immunoliposome during immunochemotherapy is released in intracellular space, it has also the advantage of overcoming multidrug resistance. Further, immunoliposome may provide an ideal solution that enables the effective killing of a heterogeneous tumor by deep penetrating the tumor tissue. Immunoliposomes can exert the bystander effect, attributed to the non-targeted passive cellular uptake of drugs, implying that former internalized drugs can efflux from antigen-positive (Ag^+^) tumor cells and diffuse into and damage bystander or neighboring tumor cells ^13^. Thanks to the bystander effect, the tumor cells that are farther away from the microvessels or do not express the target antigen (i.e., antigen-negative (Ag^-^) cells) can receive a higher concentration of anticancer agents ^14^.

Tumor-induced angiogenesis is a complicated multi-scale phenomenon that includes the growth of new vessels from pre-existing vasculatures to pave the way for further tumor progression ^15, 16^. Such a neovascularization may be inspired by some chemical factors released from the tumor cells, such as vascular endothelial growth factor (VEGF) ^16^. Several pieces of evidence have proved the intensity of angiogenesis can predict the probability of metastasis in human tumors ^17^. Microvascular density (MVD), a criterion for quantifying angiogenesis, has shown prognostic value in different cancers, for instance, breast and brain ones ^17^. In summary, tumor vasculature and the process of angiogenesis form an integrated environment, in which the structures of microvessels are highly patient-specific. Solid tumors with different MVDs may also show different treatment outcomes in clinical studies. Therefore, considering different solid tumors with their personalized microenvironment (e.g., different stages of tumor growth and angiogenesis) would improve our understanding of patient care and the different therapies needed. In addition, it is possible to predict, for tumors with different vascular architectures, the success of chemotherapy treatment, which plays a crucial role in further decision-making planning by clinicians.

Various computational models have been developed to predict the treatment efficacy of classical chemotherapy as well as nano-based or targeted drug delivery systems to solid tumors ^7, 18-25^. Stylianopoulos et al. ^18^ utilized a mathematical model to examine the effect of drug parameters on drug distribution as well as the efficacy of NPs. They also investigated two multi-stage nano-drug delivery systems in a two-dimensional tumor network. However, they did not consider the heterogeneous distribution of vessel diameters, hematocrit considerations, and blood viscosity for the formation of microvascular shunts. In the following, their model was further developed by Kashkooli et al. ^7^ to study the treatment efficacy of chemotherapy and two- and three-stages of nano-drug delivery systems in a real two-dimensional tumor microvascular network. In addition, Zhan and Wang ^19^ used a multi-physics mathematical model to measure the convection-enhanced delivery of liposome encapsulated Dox in a realistic three-dimensional brain tumor. Although these models overwhelmingly focused on nano-based drug delivery through a multi-stage approach or on optimization of treatments. However, all these aforementioned studies didn’t simulate processes involved in immunochemotherapy (e.g., target-mediated drug disposition and bystander-killing effect). In addition, since the transport of payload relies more on convection and diffusion mechanisms, these two transport mechanisms are greatly dependent on the vascular structures and components, in the particular vascular system and tissue ECM ^26^. Furthermore, according to Zhuang et al. ^27^, the impact of tumor-induced angiogenesis on tumor growth can potentially affect the accuracy of fluid flow modeling. Therefore, using different vascular structures in these studies ^7, 19^ may cause the final results to depend merely on the selected tumor microvascular network.

Vasalou et al. ^20^ presented a modeling framework to simulate the effect of receptor dynamics and controllable design parameters on tumor mass shrinkage upon antibody-drug conjugate (ADC) administration through an *in vivo* study on homogeneous mouse xenografts. However, they did not consider the convective transport of ADC therapeutic agents through interstitial fluid flow. Additionally, they assumed that the cargo could not re-enter the tumor cells (i.e., no bystander-killing effect). Shah et al. ^21^ established a pharmacokinetic-pharmacodynamic (PK/PD) model to obtain comparable efficacy of ADCs parameters from *in vitro* and *in vivo* experimental systems, utilizing a kinetic cell cytotoxicity assay. This mathematical model assumed that the tissue compartment is well mixed with regard to ADC concentration and did not cover the impacts of vessel geometry and the possibility of steep gradients in the concentration of ADC therapeutic agents. Byun and Jung ^22^ developed a simplified mathematical model to take the bystander effect into account to investigate the ratio of efflux to influx, the concentration of released payload, and the proportion of Ag^+^ cells on the efficacy of ADCs. In their model, the bystander-killing effect has changed by varying amounts of target tumor cells using a fraction parameter that determines the proportion of tumor cells. A drawback of their study is that the ADC dynamics (e.g., drug-target interaction and drug internalization process) and cleaving extracellularly were not considered, nor was the direct payload diffusivity to the target tumor cells. In addition, they used a non-heterogeneous tumor cell environment, which may greatly hinder the uniform distribution of therapeutic agents and decreases their tumoricidal potential ^8^.

Burton et al. ^23^ used a system pharmacology model to predict the transport and reaction processes of ADCs exhibiting bystander effects in a three-dimensional tumor vascular network. Despite some novelties, their model was restricted to only one tumor microvascular network and they did not examine the treatment efficacy of classical chemotherapy simultaneously. In addition, they did not model the interstitial fluid flow, the effect of elevated IFP, lymphatic system, and convective transport of ADCs in tissue as well as natural drug decay, which may lead to inaccurate predictions of drug distribution. Byun and Jung ^25^ further presented a mathematical model based on stochastic differential equations (SDEs) to study the intracellular release of the payload containing a receptor-mediated endocytosis processes of targeted drug delivery systems. However, they assumed a simplified TME in which the architecture of tumor-induced angiogenesis, interstitial space of the tumor, and lymphatic drainage were not considered. Furthermore, the key processes in drug delivery, including the convection and diffusion transport mechanisms, their extravasation from blood vessels into the interstitium and the adjacent healthy tissue, as well as natural elimination due to the metabolism and physical degradation were not covered. Khera et al. ^24^ quantitatively measured the clinical impact of the bystander effect in human tumor xenograft using the PK/PD modeling approach. These computational models have, nonetheless, improved our understanding of the transport and kinetics of targeted drug delivery and supported clinical and theoretical decision-making in drug development.

There are several research gaps in the literature regarding the accurate modeling of TME heterogeneity (e.g., considering interstitial fluid flow, different microvascular networks, and lymphatic system) as well as understanding the key process of targeted drug delivery during immunochemotherapy. With this motivation, this study aims to develop a robust and comprehensive computational framework for examining the treatment efficacy of intravenous injection of Dox and immunoliposome at different stages of vascular solid tumors. First, three capillary networks with different MVDs are generated using discrete mathematical modeling of tumor-induced angiogenesis. Then, the intravascular blood flow through capillary networks is solved to find the interstitial fluid fields in both healthy and tumor tissues. Subsequently, a multi-scale mathematical model is proposed to calculate the spatiotemporal concentration of free Dox as well as immunoliposomes exhibiting bystander-killing effect during classical chemotherapy and immunochemotherapy, respectively. Ultimately, the cytotoxic assay of these two drug delivery systems (using a fraction of killed cancer cells parameter) at different stages of tumor vascularization is calculated and compared with each other.

## II. METHODOLOGY

A summary of the steps for building the present comprehensive model, as indicated in Fig. S1, is as following:

I. Generating three stages of vasculature networks with different MVDs;
II. Calculating vascular blood flow through capillary networks with adaptable microvessels and non-continuous behavior of blood;
III. Solving mass and momentum conservation equations to obtain interstitial fluid flow parameters within both healthy and tumor tissues;
IV. Solving mass transport to obtain the different drug concentrations in intracellular and extracellular spaces during classical chemotherapy as well as immunochemotherapy; and
V. Cytotoxicity assay through calculating the fraction of killed cancer cells parameter.

### A. Description of mathematical models

#### 1. Tumor-induced angiogenesis, vascular blood flow, and microvessel adaptation

A detailed description of the mathematical modeling of tumor-induced angiogenesis and related parameter values are presented in the supplementary file and our previous work ^28^.

#### 2. Interstitial fluid fields

The tumor and adjacent healthy tissues can be assumed to be a porous environment ^28^. The interstitial fluid parameter is determined by coupling the conservation equations for momentum and mass. The momentum conservation equation for incompressible and Newtonian fluid within a porous environment (by neglecting the friction within the fluid as well as the solid and fluid stages) is simplified to Darcy’s law at a steady-state phase ^28^. Darcy equation, which describes the convective contribution of the interstitial fluid fields, is given as follows ^28^:

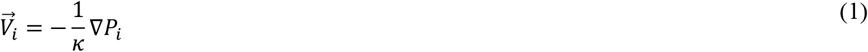

where 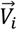 is the interstitial fluid velocity (IFV), *P*_*i*_ is the IFP, and *κ* is the hydraulic conductivity of the interstitial fluid.

The mass continuity equation for interstitium considering the presence of source and sink terms of mass in biological tissues is corrected as ^28^:

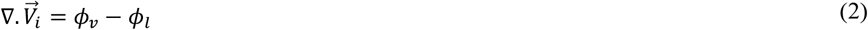

where *ϕ*_*v*_ is the rate of fluid flow from the microvascular network to the ECM and *ϕ*_*l*_ is the rate of fluid flow from the interstitial space to lymph vessels, which is calculated using Starling’s law as ^28^:

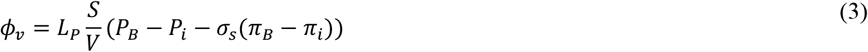

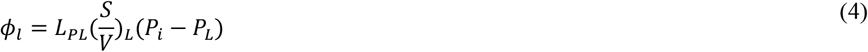

in which ^*s*^/_*v*_ is the surface area of the blood microvessels per unit volume of tissue, 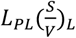 is the coefficient of lymphatic filtration, and *P*_*L*_ is the lymphatic pressure. The corresponding parameter values along with their definitions are presented in Table SI.

#### 3. Classical chemotherapy (free Dox delivery)

The transport of drugs is described by PDEs to evaluate the spatiotemporal distribution of anticancer drugs within the tumor tissue. This series of equations is also known as convection-diffusion-reaction (CDR) equations.

The key assumptions considered in the present comprehensive model for the Dox transport are as the following:

- Transport across microvessels through convection and diffusion mechanisms;
- Transport through the ECM by diffusion and convection mechanisms; and
- Binding to tumor cells, degradation through ECM, as well as cellular internalization.

For the transport of Dox during classical chemotherapy, the CDR equations are modified as ^7^:

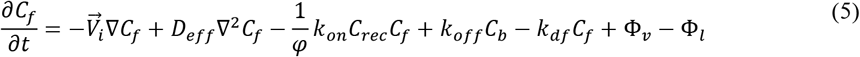

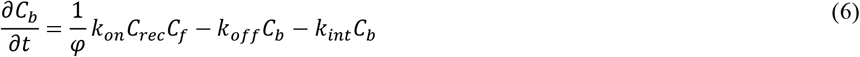

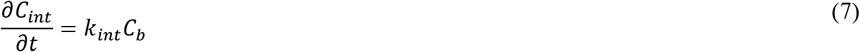

in which the free Dox (*C*_*f*_) can bind and/or unbind to tumor cell receptors with *k*_*on*_ and *k* _*off*_ constant rates, respectively, and/or it may degrade at the interstitial space by a constant rate of *k* _*d f*_. Bound drug (*C*_*b*_) can be internalized to the tumor cells by a constant cellular uptake rate of (*k* _*int*_) to be internalized drug (*C*_*int*_). Dox transport parameters along with their definitions and values are listed in Table SII.

#### 4. Targeted nano-sized drug delivery system (immunochemotherapy)

In this study, essential biological phenomena regarding immunoliposome transport are considered, as demonstrated in Fig. 1, as:

- Immunoliposome transport across microvessels through convection and diffusion mechanisms;
- Transport through the ECM through diffusion and convection mechanisms;
- Release of Dox drugs in the interstitial space due to instability of immunoliposomes;
- Binding to antigen-positive tumor cells, internalization and releasing intracellularly; and
- Bystander-killing effect following degradation of free Dox within the ECM.

**FIG. 1.**
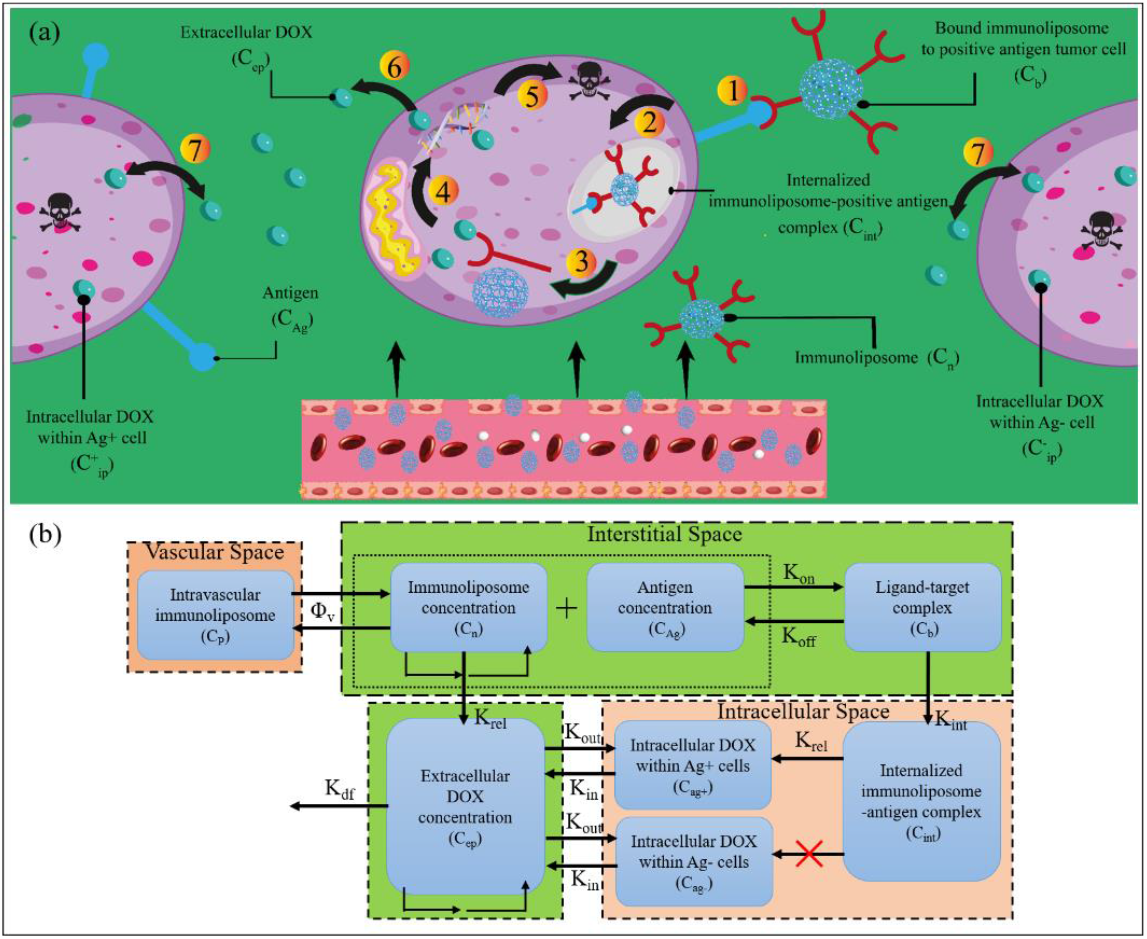
Framework of sequential steps for transport of immunoliposomes and the corresponding multi-compartment model used in the mathematical modeling. **(a)** After intravenous injection of immunoliposomes, they are carried into the tumor via the bloodstream and then cross the gaps between endothelial cells to the interstitial space. After the extravasation of the immunoliposomes, they penetrate the tumor via convection and diffusion transport mechanisms. (1) Free immunoliposomes may bind or unbind to the cells with specific antigen receptors (Ag^+^ tumor cells). Additionally, it is assumed that these free immunoliposomes may be degradated, due to natural elimination or be out of access to tumor cells at interstitial space, which causes Dox to be released; (2) Immunoliposome and its target antigen are internalized by Ag^+^ tumor cells; and then (3) They penetrate to the lysosome where the immunoliposomes are cleaved to release their anticancer cargo; (4) Due to the high toxicity of intracellular Dox, sufficient accumulation of them within the tumor cells may damage DNA; which ultimately result in (5) Cell death; (6) Due to the bystander effect, free extracellular Dox molecules can also diffuse out of Ag^+^ tumor cells; and (7) Penetrate neighboring cells (Ag^+^/Ag^-^ tumor cells) to cause cell death and/or diffuse through the interstitial space. **(b)** Immunoliposome transport multicompartmental modeling of current work (*Φ*_*v*_: exchange of therapeutic agents between capillary network and interstitium, *k* _*on*_: immunoliposome binding rate to antigen, *k* _*off*_: immunoliposome-antigen dissociation rate, *k* _*int*_: internalization rate constant, *k* _*rel*_: release rate constant, *k* _*out*_: free payload uptake rate into cells, *k* _*out*_: free payload efflux rate from cells, *k* _*d f*_: degradation rate constant).

For the transport of immunoliposomes during targeted nano-sized drug delivery system, the system of CDR equations is adjusted thus ^7, 18, 23^:

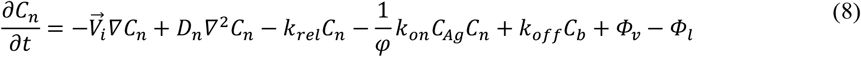

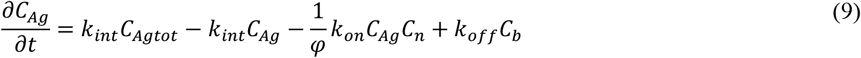

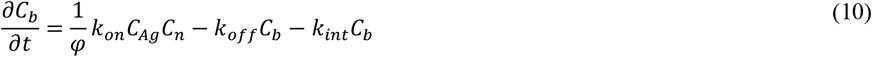

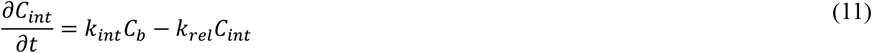

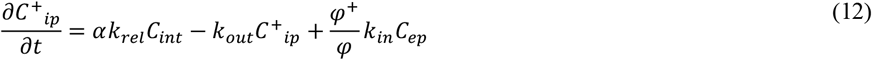

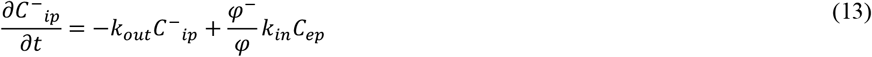

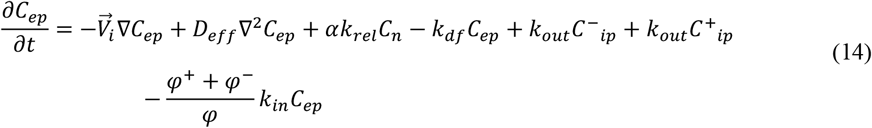

in which *C*_*n*_ is the immunoliposome concentration, *C*_*Ag*_ is the antigen concentration, *C*_*b*_ is the concentration of bound immunoliposome to positive antigen tumor cells (ligand-target complex), *C*_*Agtot*_ is the sum of *C*_*Ag*_ and *C*_*b*_ as the total antigen concentration, *C*_*int*_ is internalized immunoliposome-positive antigen complex concentration, *C*^+^_*ip*_ is intracellular Dox concentration within Ag^+^ cells, *C*^−^_*ip*_ is intracellular Dox concentration within Ag^-^ cells, and *C*_*ep*_ is extracellular Dox concentration. *φ* is the tumor volume fraction available for the drugs, *φ*^+^ is the fraction of antigen-positive tumor cells, and *φ*^−^ is the fraction of antigen-negative tumor cells. The related parameters used for immunoliposome transport with their definitions are listed in Table SIII.

In Eqs. 5 and 8, *Φ*_*v*_ is the rate of drug transport per unit volume from the blood vessels into the interstitium and *Φ*_*l*_ is the drug transport rate per unit volume from interstitial space into the lymph system. They are defined as ^28^:

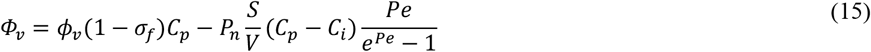

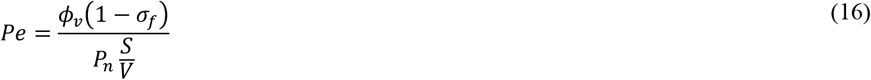

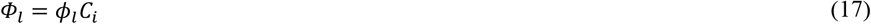

in which, *C*_*p*_ is the injected drug concentration as defined in Eq. 18, *σ*_*f*_ represents the filtration reflection coefficient, *P*_*n*_ is the permeability coefficient of microvessels, and *Pe* is the Peclet number. Regarding classical chemotherapy and the immunoliposome drug delivery system, *C*_*i*_ is attributed to the Dox concentration (*C*_*f*_) the immunoliposome concentration (*C*_*n*_), respectively.

It should be noted that during both classical chemotherapy and immunoliposome therapy, a bolus injection of drugs, indicating the vascular concentration, is considered as ^7, 18^:

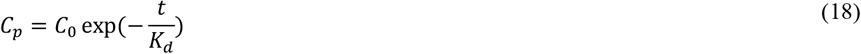

where *C*_0_ is the initial concentration and *K*_*d*_ is a time constant representing the half-life of the drug in blood circulation.

#### 5. Cytotoxicity assay and microvascular density calculation

Dox is a well-known and popular chemotherapeutic agent due to its efficacy in damaging a broad range of tumors (e.g., carcinoma, lung, and breast cancers), which generally damage the DNA of tumor cells. Employing the internalized drug concentration, the anticancer effects of drug agents can be determined by the fraction of killed cells (FKCs) metric using an empirical equation. FKCs for Dox anticancer drug is determined as ^7, 18^:

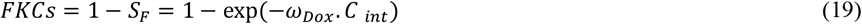

in which *S*_*F*_ is the fraction of survived tumor cells after therapy and *ω*_*Dox*_ is a constant determined for Dox according to the experimental data ^29^ and *C* _*int*_ is the drug concentration in intercellular space.

The AUC, a commonly used pharmacokinetic parameter, indicates the actual extracellular drug amount in the interstitium available to the tumor over time, which can be described as ^7^:

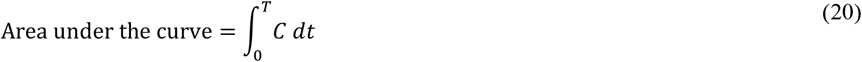

in which T is the period of treatment and *C* is the drug concentration in the extracellular space. To obtain quantitative representation and additional insight into the dynamic changes taking place in the microvascular architecture over different tumor networks, a new parameter is introduced by ^30, 31^:

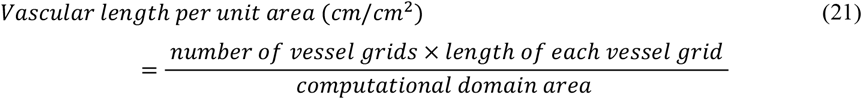

### B. Boundary conditions

A 2D equilateral rectangular computational domain is modeled in which a circular solid tumor is located at the center of its adjacent healthy tissue (Figure S2). Further, the capillary networks start to grow from two parent vessels on the vertical lines of the domain to achieve a more realistic TME. To examine the effect of MVD and angiogenesis rate on the drug delivery system, three different microvascular networks with their unique architecture in a 2-cm tumor are generated. These microvascular networks are selected according to the previous studies so that the tumor-induced angiogenesis process has been initiated ^32-34^.

For calculating blood flow through microvessels, the inlet and outlet pressure values for both parent vessels have been chosen based on the physiologically-accurate boundary conditions determined in the literature ^35^ as:

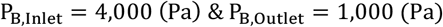

A continuity boundary condition for interstitial fluid flow and different concentrations as well as their flux are selected at the inner boundary (i.e., the boundary between tumor and healthy tissues). Different boundary conditions utilized in the interstitial fluid flow are outlined in Table SIV, where Ω_*n*_and Ω^*t*^ denote the healthy and tumor tissues at their interface, respectively. Since the IFP at the outer boundary (i.e., boundary around healthy tissue) is fixed, a Dirichlet boundary condition is used ^28^. For the transport of therapeutic agents (i.e., classical chemotherapy and immunochemotherapy) at the outer boundary, an open-type boundary condition is used ^28^.

### C. Solution strategy and computational modeling

Present model includes four important parts: (1) generating capillary networks induced by solid tumors, (2) solving interstitial fluid flow equations, (3) solving CDR equations for classical chemotherapy, and (4) solving CDR equations for immunochemotherapy. A block diagram showing different phases of modeling for each part is shown in Fig. S1 to clarify the computational techniques involved in this work.

The primary part of the model is tumor-induced angiogenesis, which contains biochemical agents, including ECM (fibronectin gradient-induced haptotaxis for sprouting), VEGF gradient-induced chemotaxis, and endothelial cell (EC) phenotypes (tip cell migration). The model also incorporates microvessel growth and remodeling, which is affected by mechanobiological and biochemical signals from wall shear stress with precise hemodynamics and hemorheology. The whole procedure for the first part of the model is as follows:

1. Assuming initial values for intravascular blood pressure (IBP), hematocrit (H_D_), blood viscosity (μ_app_), and diameter of microvessels (D_v_);
2. Calculating IBP iteratively according to governing supplementary Eqs. 6-17;
3. Updating blood viscosity at each vessel and obtaining blood hematocrit according to Pries et al. ^36^;
4. Calculating metabolic stimuli and blood hemodynamic according to vascular remodeling. See ^28, 30^ for details;
5. Updating vessel diameters based on the new values calculated in step 5;
6. Calculating the maximum relative error for IBP and D_v_ by Max(X^N^ – X^N-1^)/X^N-1^ and checking with a 10^−6^ threshold solution;
7. Reasonable IBP values are then used to calculate the interstitial fluid fields and drug transport.

The computational domain for different microvascular networks of tumors is selected to be 5 cm ×5 cm with 200×200 lattice sites in the domain in which the length of each microvessel lattice grid is 10 μm. The lattice-based grid generation is used to decrease both time and computational costs. The ECs are assumed to move one step in the tissue during each time step of the simulation. The time step of the simulation is selected in such a way that the EC movement speed in the tissue is 14 μm/h ^30^. The governing equations related to tumor-induced capillary growth are discretized using Euler finite difference method. Due to the complexity of angiogenesis mathematical modeling, a computational code is developed to solve the angiogenesis procedure and update the capillary diameters and IBP calculations. Based on the superior object-oriented ability of this programming language, vessels and nodes are considered distinct objects with appropriate features attributed to them. In addition, since the calculation of blood flow in microvessels includes a set of non-linear equations, an iterative successive over-relaxation (SOR) algorithm is proposed to calculate IBP at each node of the capillary networks. Ultimately, three microvascular networks are generated based on the different MVDs.

Subsequently, in the second part of the model, the mass and momentum equations in the interstitial fluid flow (Eqs. 1-4) are solved in a steady-state phase, after which the architecture of capillary networks is qualitatively verified. IFP and IFV parameters are then used to solve time-dependent CDR equations to obtain the spatiotemporal distribution of different therapeutic agents. To solve them, an iterative process is used. The solution starts with an initial value for different therapeutic concentrations and then a new value is calculated in each time step. This value is used as an updated value at the next time step and this process continues until the specified final time. In the case of classical chemotherapy, the final time is 10 h and for immunochemotherapy is 250 h. Ultimately, when the different intracellular, extracellular, and internalized spatiotemporal distribution of drugs are obtained, the treatment efficacy of classical chemotherapy and immunochemotherapy is also investigated by calculating the FKCs and AUC in each tumor network according to Eqs. 19 and 20, respectively.

The governing equations for fluid flow and transport of therapeutic agents, including continuity, Darcy, and CDR equations are solved using the commercial CFD software COMSOL Multiphysics 5.5 (COMSOL, Inc., Burlington, MA, USA), which works based on the finite element method. Four orders of magnitudes are chosen for the residual square errors. Multifrontal massively parallel sparse direct solver (MUMPS) is considered the direct solver with the backward differentiation formula (BDF) time-stepping method. A time step of 10 s is used; smaller time steps have not significantly altered the results but increased the computational time. The grid independency test is also performed and the results of IFP and intracellular concentration of drugs for four different computational grids (i.e., coarse, medium, fine, and extremely fine) are evaluated. There is a maximum of lower than 3% difference between the medium and fine grids, and lower than 1% variation between the fine and extremely fine grids. Therefore, due to the lowest computational costs, the fine grid is considered and used in the subsequent simulations. The fine grid has about 44,350 triangular elements, of which 660 elements are on the edges and the average element quality of the grid is 0.92. All the computational simulations are carried out on a personal laptop with CPU Intel (R) Core i5-6200U processor, 2.4 GHz CPU, and 8 GB memory.

### D. Model validation

The accuracy and validity of the present computational model are successfully confirmed by comparison with numerical results and experimental data of the previously-published studies. Since the present model has consisted of different equations, including the parabolic equations for tumor-induced angiogenesis, Hagen-Poiseuille, Darcy, and CDR equations, it is necessary to evaluate the accuracy of each component.

A qualitative comparison between the three microvascular networks considered in the present study and biological data of solid tumor morphology is made. The microvascular networks generated by our mathematical modeling are in good agreement with the *in vivo* observations ^37-39^, in which the branching of new vessels increases nearby the tumor periphery as well as within the tumor domain, where the gradient of tumor angiogenetic factors is relatively higher than that of the surrounding healthy tissue. How the microvessels grow from their initial capillary sprouts emerging from the parent vessel is also qualitatively validated by the *in vivo* tumor-induced capillary architectures. In addition, the architecture of our tumor networks with different MVDs through the TME captures the evolution of a real pre- and post-antiangiogenesis network of tumor-penetrating neo-vessels ^37-39^. Furthermore, according to Fig. 3a, the average value of measured IBP within the microvascular networks is in excellent agreement with the reported physiological values (i.e., 2,660 to 3,333 Pa) ^40^. Additionally, this measured IBP has an acceptable consistency compared to a recent numerical study of d’Esposito et al. ^41^, in which they investigated that the average vascular pressure in the LS174T human colorectal carcinoma tumor is 27,799.7 Pa.

Comparisons of our IFP and IFV results with experimental measurements of Boucher et al. ^42^ as well as numerical predictions of Soltani and Chen ^43^ and Al-Zu’bi and Mohan ^44^ are shown in Figs. 2a and 2b. The average spatial value of IFP in the tumor equals 1,359.96 Pa. The mean IFP in the healthy tissue is obtained at roughly 50 Pa, which is considerably lower than the tumor IFP. Additionally, there is a sharp IFP gradient (and consequently, a high IFV field) at the interface boundary between the tumor and healthy tissues, as illustrated in Fig. 2b. The trend of these results and their magnitudes are in excellent agreement with the previous numerical studies ^44, 45^ as well as experimental data in the literature ^42^. In addition, as shown in Figs. 3a and 3b, the measured mean IFP in the present model has high compatibility with previous numerical studies ^28, 30^ and experimental observation ^46^, which have reported about 1,533.32 Pa for IFP inside the tumor. The solid tumors modeled by Al-Zu’bi and Mohan ^44^ and Soltani and Chen ^43^ were avascular, while the current model is based on vascularized tumors; therefore, our model’s IFP value is relatively higher. In addition, the present model shows an excellent agreement for the order of IFV magnitudes with the previous numerical studies ^44, 45^. Further, a remarkable qualitative agreement is found between the IFP and IFV values obtained by our model and the corresponding parameters obtained using *in vivo* ultrasound poroelastography tests by Islam et al. ^47^. The distribution of IFV can be predicted by the Darcy equation (as the IFV value is merely proportional to the IFP gradient) and as a result of the extreme drops in the IFP value over the tumor borders. This issue is in great agreement with the experimental study of Butler et al. ^48^ and Islam et al. ^47^ as well as several numerical studies ^28, 30^.

**FIG. 2.**
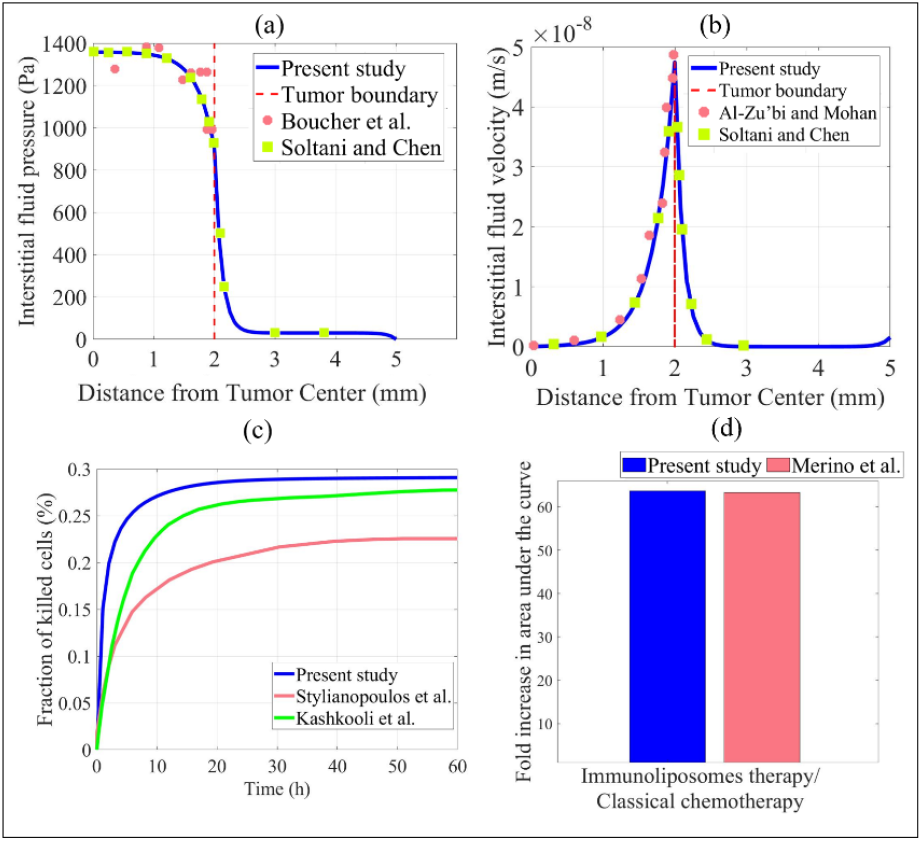
Accuracy and validity of the present model compared with numerical and experimental studies. Comparison of **(a)** interstitial fluid pressure (IFP); and **(b)** interstitial fluid velocity (IFV) fields in the tumor and healthy tissues with the radial distance from the tumor center with experimental data of Boucher et al. ^42^ as well as numerical studies of Soltani and Chen ^43^ and Al-Zu’bi and Mohan ^44^. **(c)** Comparison of fraction of killed cells (FKCs) value 60-hour post-injection classical chemotherapy obtained from the experimental data by Stylianopolous et al. ^18^ and numerical study of drug delivery in a real tumor image by Kashkooli et al. ^7^ with our model. **(d)** Comparison of the fold increase in AUC (i.e., drug concentrations over time) in tumor tissue between the present computational model and the *in vivo* study of Merino et al. ^49^.

**FIG. 3.**
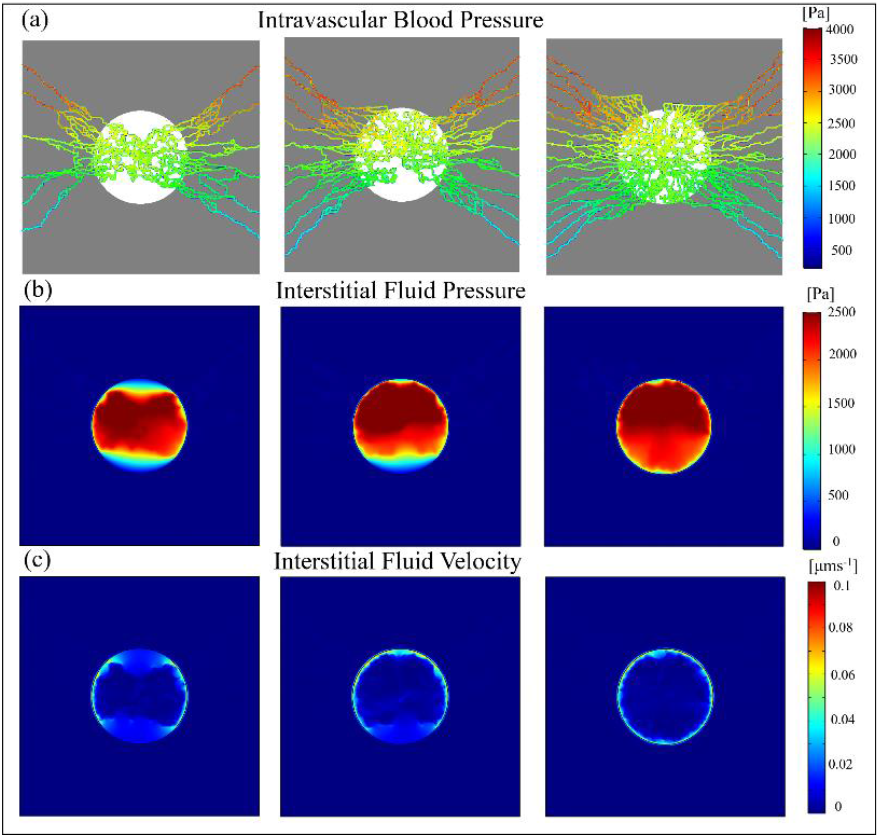
Intravascular and interstitial fluid fields. Distribution of **(a)** intravascular blood pressure (IBP), **(b)** interstitial fluid pressure (IFP), and **(c)** interstitial fluid velocity (IFV) in three generated microvascular networks (first column: low MVD, second column: intermediate MVD, third column: high MVD).

CDR equations employed in the present drug delivery systems are similar to the equations of drug transport by Stylianopoulos et al. ^18^ and Kashkooli et al. ^7^. Stylianopoulos et al. ^18^ have successfully validated their model compared to *in vivo* data for the delivery of anticancer drugs in murine mammary carcinomas. Therefore, to examine the accuracy of the present drug delivery system, the same parameter values of chemotherapy are applied, as listed in Table SII. Figure 2c compares the mean spatial FKCs in the tumor tissue over time, indicating the predicted FKCs have an acceptable agreement with the results of Stylianopolous et al. ^18^. In addition, our results show a remarkable correspondence with the FKCs values obtained by Kashkooli et al. ^7^, which investigated the treatment efficacy of classical chemotherapy in a real image extracted from mice tumor containing capillary network. It is worth mentioning that the minor discrepancy in FKCs values might be referred to differences in the microvascular networks, tumor shape, and computational domain dimension.

Since measuring drug concentrations in preclinical *in vivo* models is very complicated, suitable experimental results are not available to validate the mathematical models. Accordingly, the accuracy of the targeted nano-sized drug delivery equations to calculate immunoliposome distribution during immunochemotherapy has been mainly validated against an experimental investigation ^49^ qualitatively. Figure 2d shows the comparison of the fold increase in AUC of drug concentrations over time in tumors between the present computational model and the *in vivo* experiments of Merino et al. ^49^. These results have a remarkable agreement; in which the mean-percentage difference between our mathematical modeling and *in vivo* tests is about 0.64%, indicating that the results of the present study are sufficiently reliable for further investigations.

A qualitative comparison of the local distribution of free Dox in our model and representative three-color composite distribution of Dox *in vivo* is also made. In both *in vivo* measurements by Primeau et al. ^50^ and our computational simulations, the concentration of intracellular Dox is mainly spread heterogeneously throughout the networks with markedly raised values around the periphery of blood microvessels.

## III. RESULTS

### A. Tumor-induced angiogenesis with different microvessel densities

The recruitment of new blood vessels from adjacent vasculature during the angiogenesis process is an important factor in tumor growth, which influences the further expansion of the tumor and its response to systemic therapies. The present work is based on a study of vascular networks generated by a discrete mathematical model of tumor-induced angiogenesis, which accounts for the formation of a capillary network dependent on chemical stimuli released by the solid tumor.

To investigate the various stages of tumor vascularization, days 16, 48, 81, 121, and 145, in which the growth of microvessels initiated from the parent vessel, are selected (Fig. S3). At day 16, VEGF released by the solid tumor produces a gradient that extends the primary vessel and leads to sprouting angiogenesis to create new microvessels. In the present study, to generate microvascular networks with different densities, two parent vessels with 5, 10, and 15 initial sprouts on each one have been chosen for the low, moderate, and high MVD networks, respectively. The new microvessels elongate, branch, deform and extend toward the solid tumor during days 48-81. After day 81, the transition between the avascular and the mortal vascular tumor growth phases is terminated as new capillaries penetrate the tumor tissue. During the growth of microvessels in vascular tumors (days 81-121), heterogeneities appear as a consequence of the non-uniformly generated microvessels and asymmetric distributions of tumor angiogenesis factors. As a result of the higher VEGF gradient within the tumor, the freedom of movement of ECs is increased, and hence microvessels inside the tumor tissue become more tortuous, while their movements over the healthy tissue nearby the tumor domain are restricted. Ultimately, the vascular length per unit area (cm/cm^2^) parameter, at day 145, for low, moderate, and high MVD networks is 50.15, 63.53, and 88.18, respectively. The microvascular networks at day 145 are used to investigate the delivery of therapeutic agents.

### B. Interstitial and intravascular fluid fields

After generating the microvascular networks, the IBP within the three networks is separately calculated based on Poiseuille’s law. In addition, given the biological phenomena in porous media, the interstitial fluid parameters (IFP and IFV) are described by Darcy’s law. The results of IBP, IFP, and IFV are illustrated in Fig. 3.

The average IBP in all the examined microvascular networks differs around 2,996.69 Pa with no considerable differences among each network. The present mathematical modeling does not predict significant differences between the networks with various densities of vasculature in terms of IBP. Apart from networks, the maximum value of IBP occurs near the left sides of the field (inlets), but its value moderately decreases near the right sides (outlet) due to the pressure drop.

In all the microvascular networks, the IFP has its maximum value in the tumor compared to normal tissue. As the MVD increases, the IFP through the tumor tissue is also increased. The average spatial values of tumor IFP in high, intermediate, and low MVD networks are obtained at 2,278.52, 2,116.94, and 1,958.92, respectively. In each network, the IFP has relatively higher values in some regions that have a greater MVD. Additionally, the IFP has its minimum value in the top and low tumor regions of the network with low MVD. Similarly, the same pattern in IFP results but in a limited region of the intermediate MVD network is seen, while the spatial distribution of IFP in the high MVD network is relatively more homogeneous. In other words, both the local distribution of IFP and its magnitude greatly depend on the MVD.

The IFV has a low value (in order of 10^−8^ m.s^-1^) throughout the whole domain unless at a thin layer at the interface of tumor and healthy tissues, where the maximum value occurs as a result of a large IFP gradient therein. The average IFV inside the tumor network with high MVD is 8.69×10^−9^ m.s^-1^, while this value increases to 9.74×10^−9^ and 1.02×10^−8^ m.s^-1^ in the intermediate and low MVD networks, respectively. Similar to the low magnitudes of IFP at the location with lower MVDs, IFV is also decreased, in which a complete circular pattern of elevated IFV has been created at high MVD networks.

### C. Classical chemotherapy

During classical chemotherapy, the intravenously injected drug molecules (here, Dox) are transported into the ECM by crossing the walls of blood vessels, thereafter diffusing to the tumor interstitium. Drugs can then bind or unbind to the receptors of tumor cells, degrade within the ECM, and/or internalize to the tumor cells. Internalized drugs, intracellular concentration, can eliminate the cancerous cells by their anticancer effects through DNA damaging. In the current model, the transport of cytotoxic payload is measured by CDR equations, in which the drug diffusion occurs in response to a concentration gradient, based on Fick’s first law of diffusion.

Figures 4a-c show different mean spatial concentrations of Dox during classical chemotherapy. The decay in concentration, after reaching the maximum magnitude, is a result of a decrease in the intravascular released drug, elimination through the blood microvessels and interstitial space, as well as cellular uptake. The bound Dox concentration shows the same trend for different networks but with approximately four-fold lower amplitude. 10-hour post-injection of free Dox, internalized Dox concentration reaches its maximum value within the tumor in all the examined networks. Different concentrations of the drug are increased as the MVD decreases in which the internalized drug at low MVD is ultimately increased up to 1.3 and 1.9 times higher than intermediate and high MVDs, respectively. Figures 4d-f illustrate different spatial concentrations of internalized Dox in different networks after 10-hour classical chemotherapy. Decreasing MVD results in higher internalized Dox; thereby the treatment efficacy of chemotherapy increases. However, the intracellular Dox in all the tumor microvascular networks has a limited extracellular drug penetration and principally accumulated nearby the blood vessels with a heterogeneous distribution, restricting the treatment efficacy.

**FIG. 4.**
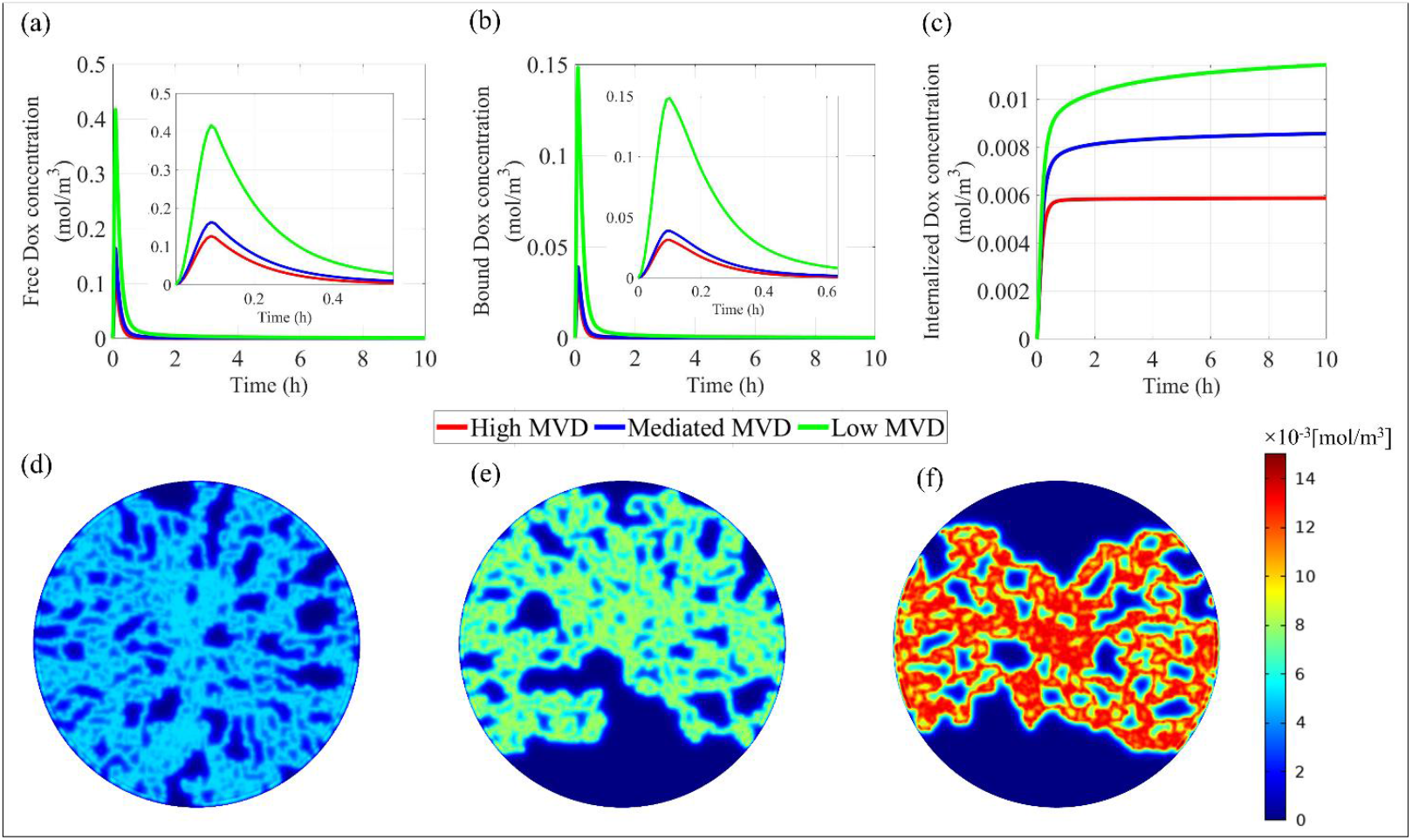
Spatiotemporal distribution of drug at different networks during classical chemotherapy. Mean Spatial-temporal concentration profile of **(a)** free, **(b)** bound, and **(c)** internalized Dox in the tumor tissue considering different microvascular networks. Spatial distribution of internalized Dox after 10-hour chemotherapy in tumor microvascular networks with **(d)** high, **(e)** intermediate, and **(f)** low MVDs.

### D. Immunochemotherapy

Figure 5 shows different temporal concentration profiles of the drug concentrations across the tumor and the corresponding treatment efficacy achieved from the delivery of immunoliposomes at different tumor microvascular networks. At the first hours, the free and bound drug concentrations increase with time due to the increase of plasma concentration and drug extravasation. Subsequently, the concentration of both free and bound drugs falls immediately to reach zero due to drug degradation and internalization phenomena. According to Fig. 5c, the concentration of internalized drug increases over time as the bound drug within the cells increases, and then because of Dox release from immunoliposomes, its value decreases gradually. As shown in Figs. 5d and 6e, the intracellular concentration of Dox within both Ag^+^ and Ag^-^ tumor cells increases slightly and ultimately reaches a constant value. After 250-h post-injection, the ratio of intracellular Dox concentration within Ag^+^ to Ag^-^ tumor cells in each network has a mean value of 2.35, corresponding to the constant rate of Ag^+^/Ag^-^ cell volume fractions 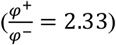. As the outward diffusion rate of tumor cells and Dox release from immunoliposomes are increased over time, the extracellular concentration of Dox within the tumor tissue at different networks increases (Fig. 5f,). In contrast, extracellular Dox concentration reduces by increasing the MVD.

**FIG. 5.**
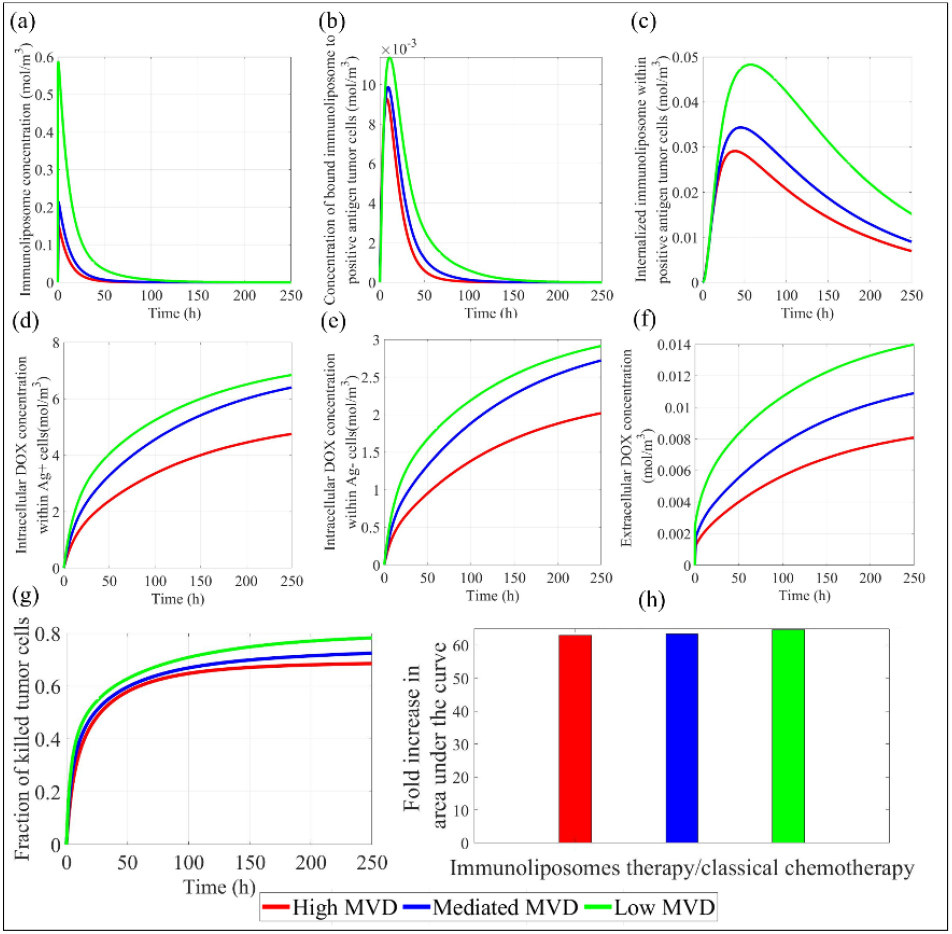
Different concentrations of drug and their response evaluations at different microvascular networks during immunochemotherapy. Mean spatial-temporal concentration profile of **(a)** immunoliposomes, **(b)** bound immunoliposomes to target cells, **(c)** internalized immunoliposomes, intracellular Dox within **(d)** Ag^+^ and **(e)** Ag^-^ tumor cells, and **(f)** extracellular Dox. **(g)** Fraction of killed cells as a function of time for different tumor networks. **(h)** Fold increase in AUC of drug concentrations over time in tumors for different networks.

**FIG. 6.**
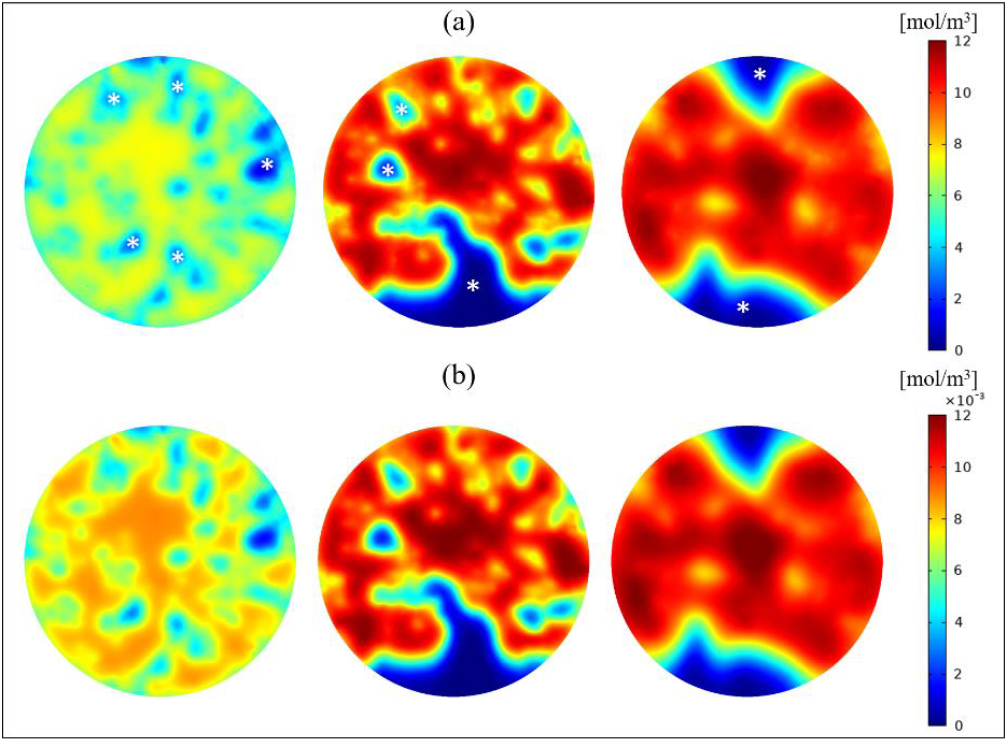
Temporal distribution of drugs in different microvascular networks during immunochemotherapy. **(a)** Total intracellular Dox within both Ag^+^ and Ag^-^ tumor cells. **(b)** Extracellular Dox concentration. (First column: high MVD, second column: intermediate MVD, and third column: low MVD).

The FKCs results shown in Fig. 5h demonstrate that the treatment efficacy of immunoliposomes enhances as the MVD of tumor networks decreases. In the first 50-h injection, owing to the lower intracellular concentration of Dox, the FKCs results among different tumor networks do not change significantly. Ultimately, 250-h post-injection, this value reaches 0.78 in low MVD networks with the highest cytotoxic activity, followed by the intermediate and high MVD networks with 7.38% and 12.40% lower than low MVD, respectively. Figure 5h summarizes that the fold increase in AUC of drug concentrations over time in tumors decreases gradually from 64.70 in low MVD network to 63.32 and 62.97 in intermediate and high MVDs, respectively. Differences in the AUC values between different stages of the tumor network are also reflected in the higher drug exposure in networks with lower MVDs expressed as FKCs amounts. To compare the drug internalization and cell interaction between classical chemotherapy and immunochemotherapy, we also calculated the AUC of the internalized drug over time within the tumor. The average of this parameter over different tumor networks during immunochemotherapy compared to chemotherapy has increased by a factor of 70.82.

The temporal distribution of the total concentration of intracellular Dox within Ag^+^ and Ag^-^ tumor cells as well as the extracellular drug is shown in Fig. 6. The distribution of intracellular drug concentration is highly dependent on the architecture and location of microvessels within the tumor as a result of the MVD-dependency of the transvascular transport. The numbers of scattered regions having low intracellular drug concertation, which have been marked by white stars, are increased as the MVD increases. As can be seen in Figs. 6a and 6b, spatial distributions of extracellular Dox compared to intracellular ones are not changed significantly at each tumor network. However, the magnitude of extracellular Dox is much lower than intracellular drugs within the tumor (almost a thousandth).

## IV. DISCUSSION

IFP and IFV are two major components of the cancer mechanical microenvironment, which play a crucial role in clinical consequences for cancer diagnosis, prognosis, and treatment ^51^. The reason for elevated IFP inside the tumor tissue in respect of healthy tissue, a hallmark of solid tumors, is originated from several biophysiological factors. The net result of high leaky blood vessels and the lack of a functional lymphatic system at the tumor tissue is interstitial hypertension thereof, thus ultimately causing intratumoral advection effects. Theoretical analyses have proved that this outward diffusion flow is the major reason for the reduction of efficacy, and heterogeneous uniformity, as well as poor transport of therapeutic and diagnostic agents into the tumor central areas ^32, 52^. Apart from these effects, interstitial hypertension increases cell clonogenicity and VEGF expression in the peritumoral tissue, which may enhance tumor-induced angiogenesis and metastasis ^8, 53^. Moreover, elevated IFV at the tumor border may also boost metastasis by rising the shear stress acting on the tumor cells and motivating them to shift into the lymphatic drainage system nearby the tumor ^54, 55^.

The decrease in vascular length per unit area in tumor networks with lower MVDs induces tumor vessel normalization by reducing tortuosity of the vasculatures and the fractional coverage of the vascular endothelium by ECs over tumor tissue. Similar striking microvascular changes have also been reported before and three days after a single injection of VEGF-receptor-2 antibody treatment by Tong et al. ^31^, in which the vascular length per unit area in their study significantly reduced from 58.8 to 41.9 cm/cm^2^. Furthermore, the results of Fig. 3b confirmed that, in constant tumor size, decreasing the MVD inside the TME leads to a reduction of IFP inside the tumor tissue. In other words, decreasing the MVD results in more normalized tumor blood vessels and lower IFP values within the tumor tissue, which ultimately causes prerequisite physiologic conditions for subsequent immuno/chemotherapy. Elevated IFP in networks with higher MVDs is attributed to the presence of dense microvasculature with complicated geometry that acts as a source term in the continuity equation (i.e., *ϕ*_*v*_ in Eq. 15). This IFP drop, as a result of decreasing in MVD, is also reported in several experimental investigations of human tumors ^56^ and transplanted tumors in mice ^31, 57^. For instance, Tong et al. ^31^ have reported that blocking VEGF signaling reduces IFP, not only by restoring the lymphatic system but also by generating a functionally and morphologically matured tumor network.

Comparing the mean spatial concentrations of the drug during the two drug delivery systems implies that immunoliposomes during immunochemotherapy have shown much larger drug internalization compared to conventional chemotherapy (Figs. 4c and 5c). This result is also reported by *in vivo* experiment of Merino et al. ^49^, in which they evaluated the combination of chemotherapy and immune checkpoint inhibitors by developing Dox immunoliposomes functionalized in a melanoma murine model. Comparing the fold-increase in internalized drug concentration within Ag^+^ and Ag^-^ tumor cells at different networks with Ag^+^/Ag^-^ cell volume fraction, similar to previous numerical and *in vivo* investigations ^58-60^, indicates that the target-antigen level is directly impacting intracellular concentration of payload. Instead, the fold-increase in internalized Dox during classical chemotherapy at different tumor networks has shown that it has nearly a linear relationship with vascular length per unit area at each network. Comparing the spatial distribution of drugs during chemotherapy and immunochemotherapy (Figs. 6, 4d-f) illustrates that the bystander-killing mechanism allows immunoliposomes to expose a larger portion of the tumor tissue to the intracellular Dox. This effect along with the intracellular release of Dox and the high capability of immunoliposomes to carry payload result in a much higher concentration of anticancer drugs with a more heterogeneous distribution compared to classical chemotherapy. Moreover, a comparison of temporal profiles of drug concentration indicates that immunoliposomes remain for a longer period in tumor networks in comparison with free Dox, leading to improvement in treatment outcomes. This result has also been reported by several numerical studies on human brain tumors ^61^. In other words, targeted liposomes encapsulating Dox have been demonstrated to be an intelligent association, in which they can selectively internalize into target tumor cells and diffuse readily from target cells, and then penetrate adjacent cells regardless of their target antigen expression. There are successfully providing a dual activity presented by both active and passive treatment mechanisms. In contrast, the net result of the heterogeneous distribution of drugs with low concentration during classical chemotherapy is that a large portion of the tumor remains untreated. This phenomenon is one of the main causes of chemotherapy failure that has also been reported in several experimental and numerical studies ^7, 18^. In other words, survived cancerous cells may regrow or enter into the circulation through fragile tumor vasculature, which is considered to be a prerequisite for metastasis ^8^.

According to Figs. 4c and 5g, regardless of implemented drug delivery systems, the intracellular concentration of the drug and its therapeutic outcomes (i.e., FKCs and AUC) increase as the MVD decreases. This result is consistent with several experimental ^31, 62^ and numerical ^30, 63^ studies. These results also provide evidence that at the initial stages of tumor-induced angiogenesis and growth, the cytotoxic agents have shown higher efficacy during chemotherapy, the phenomenon which is also reported by Vavourakis et al. ^64^. The higher cytotoxicity in tumor networks with lower MVDs strongly depends on the extent of the extravascular drug. In other words, the extravascular flow has a significant impact on the dynamics of tumor-associated vasculature growth and therapeutic response. The extent of transport of anticancer agents at interstitial space in biological tissue is determined by diffusion, motivated by concentration gradients and convection, resulting from the pressure difference between interstitial space and capillary networks ^15^. As the IFP at the networks with lower MVDs decreases, while the IBP values are relatively constant among different networks, the hydrostatic pressure gradient enhances across the microvessels. This can be found from higher values of transvascular pressure difference, ▽P=IBP-IFP, which controls the transport of drug molecules across the blood microvessel wall (Eq. 15). These findings are consistent with the experimental results of Tong et al. ^31^ and Chauhan et al. ^34^.

Although the efficacy of treatment by immunoliposomes exhibiting bystander-killing effect has been enhanced in all microvascular networks, this effect in tumors with lower MVDs has proved to act as a double-edged sword for increasing the extracellular payload (Fig. 5f). The main reason for elevated extracellular Dox concentration at networks with low MVDs should be sought in the source terms (Eq. 14). In other words, the higher concentrations of Dox within the tumor cells and free immunoliposomes in the interstitial fluid exist, the higher extracellular concentration of Dox in the extracellular space. Since the concentration of this drug type in tumor networks with a lower MVD is greater compared to the networks with a higher MVD, the extracellular payload values also increase in those networks. Although the order of extracellular payload concentration is much lower than intracellular Dox concentration, this adverse effect indicates that the selection of treatment is highly dependent on TME characteristics.

According to Fig. 6, regardless of different tumor networks, interestingly there are some tumor regions where the concentration of payloads therein is relatively lower (marked by star signs). This issue is mainly associated with the lack of blood supply, hence those tumor cells at a greater distance from microvessels would receive sub-therapeutic drug exposure. Some further factors that may affect the distribution of drugs include the intervessel distances in these cells (which are often larger than the Dox can reach them through the bystander-killing effect mechanism), the limited diffusion coefficient of Dox, and convection/diffusion barriers formed by solid tumors ^3, 61^. Similarly, Greene et al. ^65^ have reported that although the human tumor cells adjacent to microvessels would be eliminated efficiently by a specific type of ADC, those at a greater distance from tumor blood vessels would potentially escape killing. To overcome this problem, using internal (such as pH/MMP-2-sensitive nanoliposomes) or external (such as thermo-sensitive liposomes or magnetic nanoparticles activated by temperature) stimuli-responsive systems for anticancer therapies can be a promising mechanism to locally increase the drug release performance in the regions of interest ^1^. Therefore, a drug delivery system with higher clinical efficacy or combination therapies (e.g., vascular normalization and anti-angiogenic therapy or thermal ablation) is essentially needed in such stages of tumor microvascular networks.

## V. CONCLUSIONS

Present study is inspired by the performance of immunoliposomes exhibiting the bystander-killing effect, which is a smart nanocarrier for targeting the tumor cells. The treatment efficacy of such a drug delivery system has also been compared with free Dox delivery at three stages of tumor microvascular networks with different MVDs. Our results, in line with experimental studies, show that the intracellular Dox is mostly accumulated heterogeneously in the perivascular regions, owing to limited deep penetration to the tumor tissue. Furthermore, the amount of intracellular Dox concentration during immunochemotherapy is considerably increased in all the examined tumor networks compared to classical chemotherapy. This improved cytotoxicity is attributed to the bystander-killing effect exhibited by immunoliposomes and their intracellular drug release during immunochemotherapy, allowing therapeutic agents to penetrate deeper with much more concentration. However, spatiotemporal heterogeneities in blood supply and elevated IFP along with poor hydrostatic pressure gradient across the microvessel wall create an abnormal microenvironment in networks with greater MVD, impairing uniform delivery of therapeutic agents. These findings are a positive step toward treatment planning and facilitating the pre-clinical screening of antibody-based drug combinations for the future development of targeted nano-sized drug delivery systems. Additionally, this study shows how multi-scale computational models can be employed in the discovery of new anticancer drugs as well as improving our understanding of the interaction of drugs with the TME.

## SUPPLEMENTARY MATERIAL

See the supplementary material for the complete expressions of tumor-induced angiogenesis, vascular blood flow, and microvessel adaptation.

## DATA AVAILABILITY

The data that supports the findings of this study are available within the article and its supplementary material.

## AUTHORS CONTRIBUTION

Conceptualization: M.A.A. and F.M.K.; Data curation: M.A.A. and F.M.K.; Formal analysis: M.A.A.; Investigation: M.A.A. and F.M.K.; Methodology: M.A.A. and F.M.K.; Simulations and software: M.A.A.; Validation: M.A.A.; Post-processing and data visualization: M.A.A.; Writing—original draft: M.A.A.; Review and editing: M.S., F.M.K. and M.A.A.; Project administration: M.S. All authors contributed to the article and approved the submitted version.

---

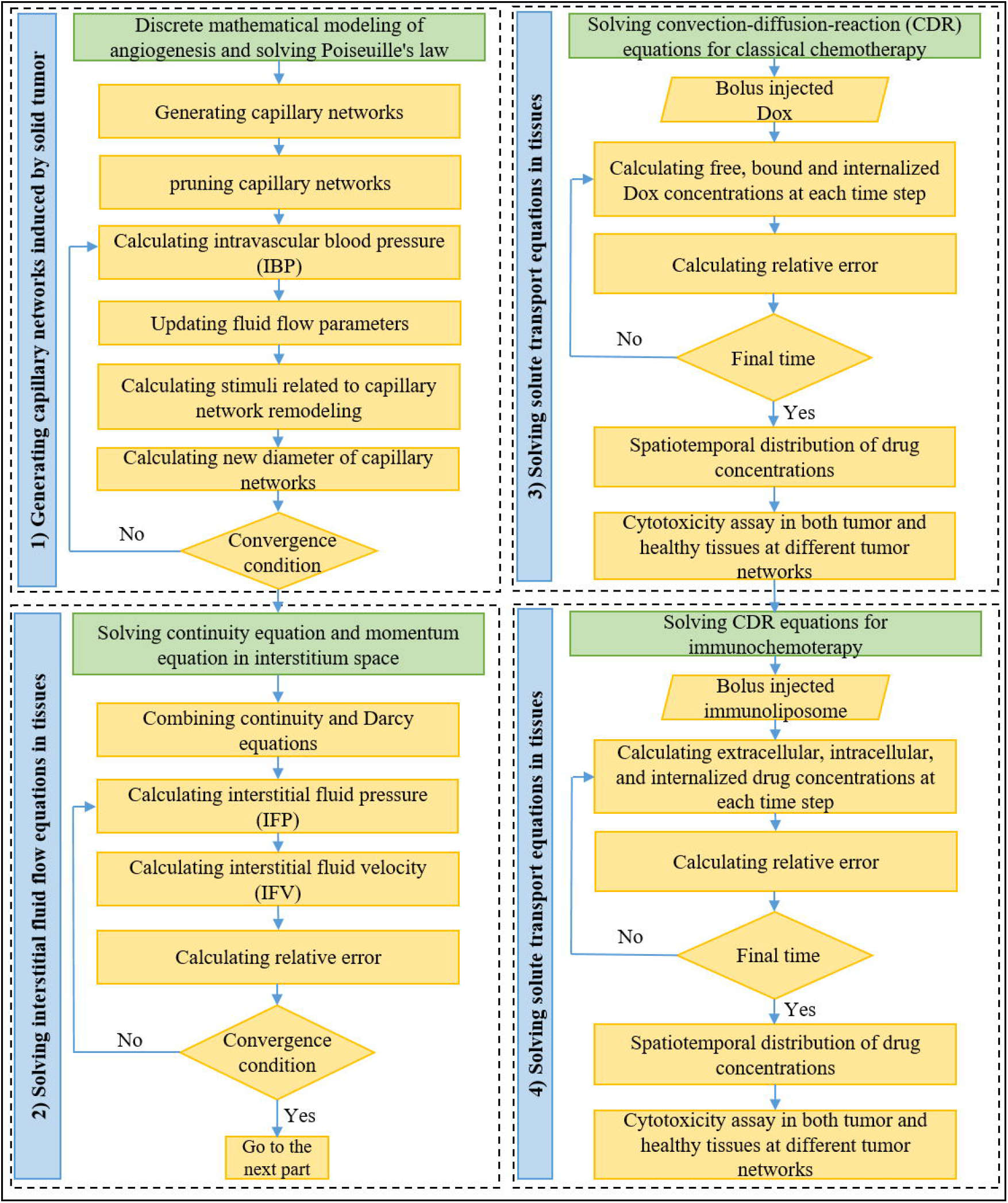

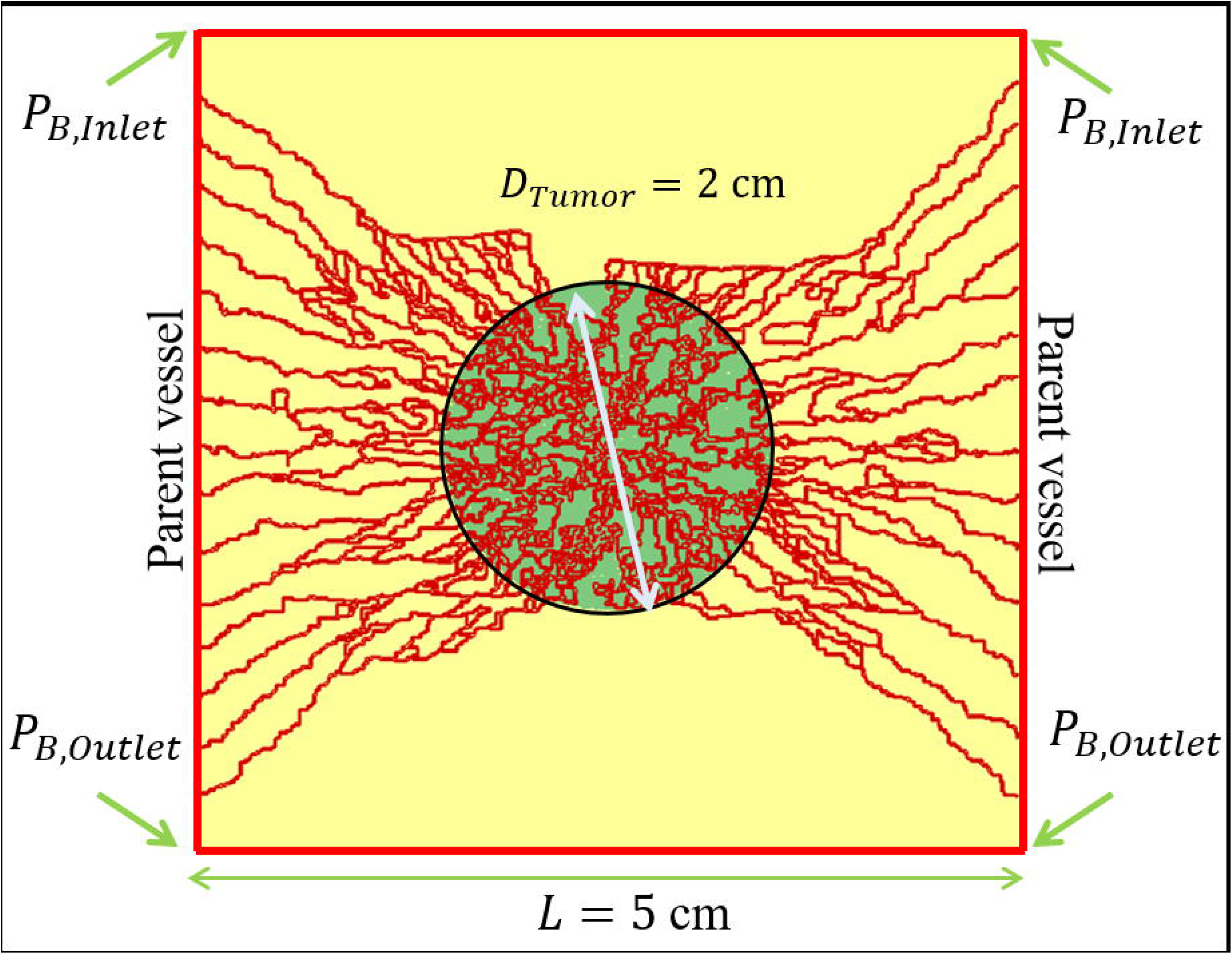

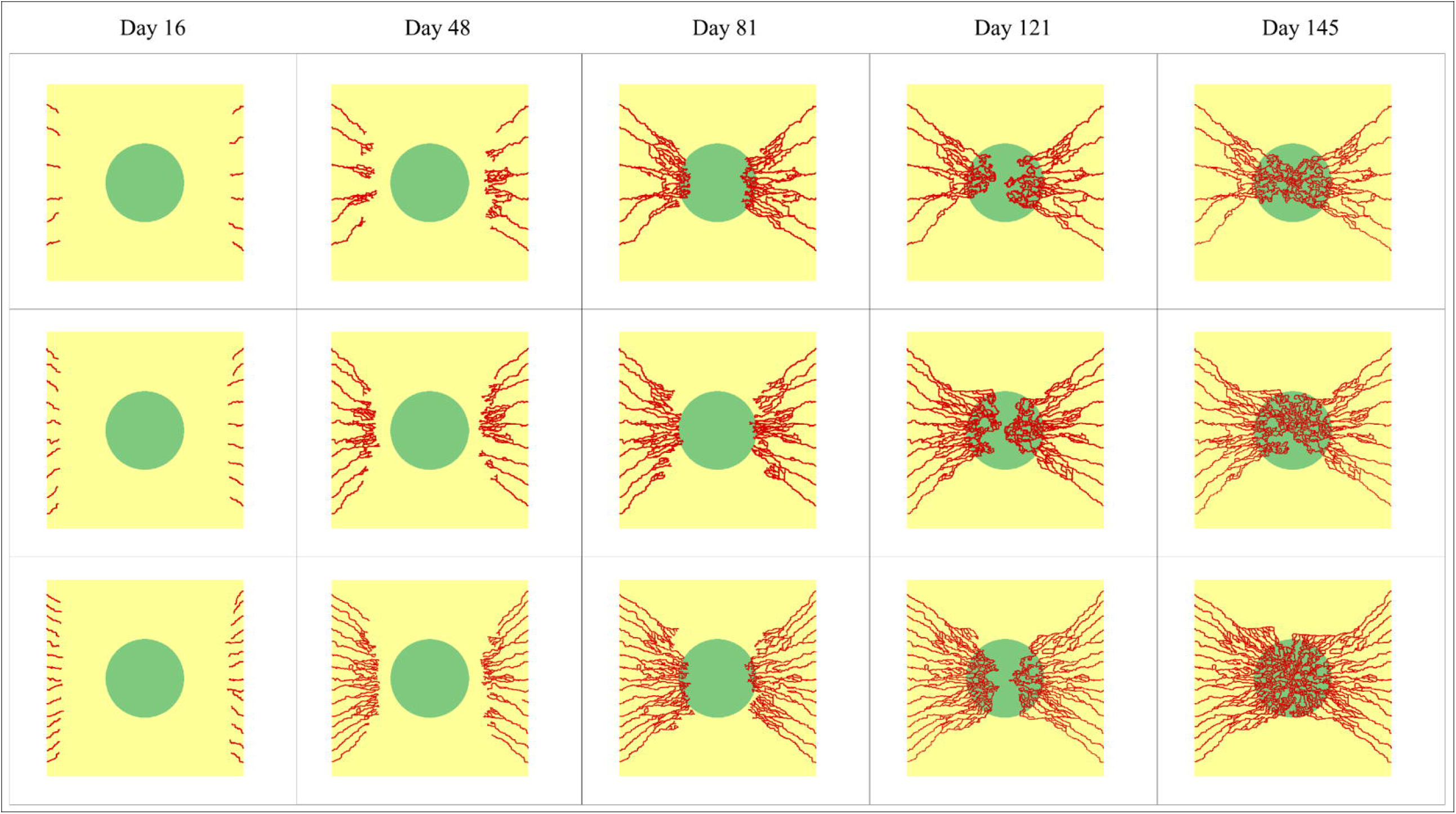

