## supplementary material for "Targeted Nano Sized Drug Delivery to Heterogeneous Solid Tumor Microvasculatures: Implications for Immunoliposomes Exhibiting Bystander Killing Effect"

### Tumor-induced angiogenesis, vascular blood flow, and microvessel adaptation

In the current study, a dynamic adaptive microvascular network modeling is proposed to simulate tumor-associated microvasculature based on a discrete probabilistic model. Such a model is initially introduced by Anderson and Chaplin [1] and then modified by our group [2, 3]. The present mathematical model captures branching, anastomosis, hematocrit, wall shear stress, and consequently blood flow induced vessel branching. The model predicts filopodias situated on the tip endothelial cells (ECs) route the trajectories and direct tip ECs, while stalk ECs proliferate and elongate the vessel [2, 3]. EC movement toward the tumor is affected by three main processes, including: (I) random movement, (II) direct movement, and (III) transverse movement. Each of the ECs movement terms can be defined with the corresponding gradient of the movement stimuli and the appropriate parameters.

The random movement is simulated like molecular diffusion as a result of the concentration gradient. Chemotaxis, which is related to tumor angiogenesis factors (TAFs) gradients secreted by hypoxic tumor cells, diffuses inside the ECM and triggers the ECs. Haptotaxis is the motion of cells due to the gradient of insoluble chemicals. Hypoxic tumor cells secrete vascular endothelial growth factor (VEGF) which is transported in the domain by diffusion. Moreover, as ECs move in the domain at the direction of the VEGF gradient, VEGF is uptaken by the ECs. Since the branching of a vessel depends upon VEGF concentration, the probability of branching surges as a vessel gets closer to the tumor. When vessels come across each other as they move in the domain, they anastomose and form closed loops [2]. The system of equations describing the combining of these three mechanisms and related formulations is represented as follows [2]:

$$\frac{\partial n}{\partial t} = D_n \nabla^2 n - R(\rho)\nabla.(\chi(c)n\nabla c) - \nabla.(\rho_0 n \nabla f) \tag{1}$$

$$\frac{\partial m}{\partial t} = \gamma n + \varepsilon \nabla^2 m - \upsilon m \tag{2}$$

$$\frac{\partial f}{\partial t} = \omega n - \xi m f \tag{3}$$

in which n is the EC density, c concentrations of VEGF, f is the fibronectin concentrations, $m$ is the concentration of chemical agent to be representative of the matrix-degrading enzyme, $D_n$ is the diffusion coefficient of EC movement, $\chi(c)$ is the chemotactic function, and, $R(\rho)$ correspond to the matrix density function. In addition, $\gamma$ and $\upsilon$ are the production and natural degradation rates of the matrix-degrading enzyme, $\varepsilon$ is the diffusion coefficient of the matrix-degrading enzyme, and $\omega$ is a constant production rate of fibronectin. The non-dimensionalized forms of the matrix density function as well as the chemotactic function, which is assumed to rise linearly with increasing the concentration of TAF, is represented below [2]:

$$R(\rho) = \exp[-\frac{(\rho - \rho_0)^2}{2\sigma^2}] \tag{4}$$

$$\chi(c) = \chi_0(1 + \chi_1 c) \tag{5}$$

where $\rho_0$ is the matrix density reference value, $\sigma$ is a constant parameter, $\chi_0$ is the chemotactic coefficient and $\chi_1$ is a positive constant.

Posterior to vasculature generation, blood flows in the network as it is a requisite to blood vessel perseverance. Based on the obtained Reynolds numbers, blood flow through microvessels is laminar [2]. Therefore, Hagen-Poiseuille's law governs the flow characteristics, as follows [2]:

$$Q_v = \frac{\pi {D_v}^4}{128 L_v} \frac{\Delta P_B}{\mu_{app}(D_v, H_D)} \tag{6}$$

where $\Delta P_B$ is the pressure difference for a microvessel segment, $L_v$ is the microvessel length, and $\mu_{app}$ is the apparent blood viscosity, which is a function of both microvessel diameter ($D_v$) and its hematocrit ($H_D$). It should be noted that the rate of change in vessel diameter is dependent upon three different stimuli, namely: wall shear stimulus, vessel transmural pressure stimulus, and metabolic stimulus [1]. The obtained intravascular blood pressure (IBP) will be used in the interstitial fluid flow model in the next subsection.

Blood contains different constituents, which make its properties variable based on the concentration of the compositions. Blood viscosity is one of the most important constituents influencing the fraction of red blood cells. Apparent blood viscosity as described by Pries et al. [4], is given as follows:

$$\mu_{app}(D_v, H_D) = \mu_{rel} \cdot \mu_{plasma} \quad (7)$$

$$\mu_{rel}(D_v, H_D) = \left[1 + (\mu_{0.45} - 1)(\frac{(1 - H_D)^c - 1}{(1 - 0.45)^c - 1})\left(\frac{D_v}{D_v - 1.1}\right)^2\right](\frac{D_v}{D_v - 1.1})^2 \quad (8)$$

$$\mu_{0.45} = 6e^{-0.085D_v} + 3.2 - 2.44e^{-0.06D_v^{0.0645}} \quad (9)$$

$$c = (0.8 + e^{-0.075D_v})\left(-1 + \frac{1}{1 + 10^{-11}(D_v{}^{12})}\right) + (\frac{1}{1 + 10^{-11}(D_v{}^{12})}) \quad (10)$$

in which $\mu_{plasma}$ is the plasma viscosity, which has a constant value of 1.2 cp, $\mu_{0.45}$ is the viscosity when red blood cell distribution is normal in the microvessel, and $c$ is a function that accounts for the shape of viscosity dependency on hematocrit.

Since the blood vessels are considered viscoelastic parts of the human body, they can dilate and/or shrink in response to the fluid that flows through them to adjust themselves to the acted shear stress. The amount of changes in the vessel diameter depends upon a total stimulus defined by Pries et al. [4]:

$$\Delta D_{\mathrm{v}} = S_{\mathrm{tot}} D_{\mathrm{v}} \Delta t \quad (11)$$

where $\Delta D_{\mathrm{v}}$ is the value of changes in diameter, $\Delta t$ is the solution time step, $D_{\mathrm{v}}$ is the initial capillary diameter and $S_{\mathrm{tot}}$ is the total stimulus. Total stimulus comprises the sum of metabolic stimulus, $S_{\mathrm{m}}$, the intravascular pressure, $S_{\mathrm{p}}$, and wall shear stress stimulus, $S_{\mathrm{wss}}$:

$$S_{tot} = S_{wss} + S_p + S_m \quad (12)$$

$$S_{wss} = \log(\tau_w + \tau_{ref}) \quad (13)$$

$$S_p = -\log \tau_e(P_B) \quad (14)$$

$$S_m = k_m \log\left(\frac{Q_{ref}}{QH_D} + 1\right) \quad (15)$$

$$\tau_w = \frac{32\mu_{app}(D_v, H_D)}{\pi D_v^3}|Q_v| \quad (16)$$

$$\tau_e(P_B) = 100 - 86.\exp(-5{,}000\ (\log(\log P_B))^{5.4}) \quad (17)$$

in which $\tau_w$ is the wall shear stress in microvessels, $\tau_{ref}$ is a constant parameter to eschew singularity at low wall shear stress rates, and $\tau_e(P_B)$ is the wall shear stress, resulting from the blood pressure. Additionally, $Q_{ref}$ , reference blood flow rate, is the rate of flow inside the parent vessel and $k_m$ is a positive constant. A detailed description of the discretization of the equations, nondimensionalization of them, initial conditions, and related parameter values are presented in our previously published work [2, 3]

**TABLE SI.** Parameters used for interstitial transport modeling.

| Parameter | Unit | Definition | Tissue type | Value | Ref. |
|---|---|---|---|---|---|
| $\pi_i$ | $mmHg$ | Oncotic pressure of interstitial fluid | Tumor | 10 | 3, 5 |
| | | | Healthy | 15 | |
| $\pi_B$ | $mmHg$ | Oncotic pressure of microvessels | Tumor | 20 | 3, 5 |
| | | | Healthy | 20 | |
| $\sigma_s$ | - | Coefficient of average osmotic reflection | Tumor | 0.91 | 3, 5 |
| | | | Healthy | 0.82 | |
| $L_P$ | $\frac{m}{Pa.s}$ | Hydraulic conductivity of the microvessel wall | Tumor | $0.36 \times 10^{-11}$ | 6, 7 |
| | | | Healthy | $2.8 \times 10^{-11}$ | |
| $L_{PL}(\frac{S}{V})_L$ | $\frac{1}{Pa.s}$ | Coefficient of lymph filtration | Healthy | $1 \times 10^{-7}$ | 3, 5 |
| $\kappa$ | $\frac{cm^2}{mmHg.s}$ | Hydraulic conductivity of interstitium | Tumor | $4.13 \times 10^{-8}$ | 3, 5 |
| | | | Healthy | $8.53 \times 10^{-9}$ | |
| $P_L$ | $Pa$ | Hydrostatic pressure of lymph vessels | Healthy | 0 | 3 |

**TABLE SII.** Parameters used for classical chemotherapy.

| Parameter | Unit | Definition | Tissue type | Value | Ref. |
|---|---|---|---|---|---|
| $D_{eff}$ | $m/s$ | Diffusion coefficient of nanocarrier | Tumor | $3.4 \times 10^{-10}$ | 8 |
| | | | Healthy | $1.58 \times 10^{-10}$ | |
| $P_n$ | $m/s$ | Microvessel permeability coefficient | Tumor | $3.00 \times 10^{-6}$ | 9 |
| | | | Healthy | $3.75 \times 10^{-7}$ | |
| $\sigma_f$ | $m^2/s$ | Reflection coefficient | - | 0.35 | 8 |
| $K_d$ | $min$ | Blood circulation decay constant | - | 6 | 8 |
| $k_{on}$ | $1/(M.s)$ | Binding rate constant | - | $150$ and $1.5 \times 10^{4*}$ | 8, 10, 11 |
| $k_{off}$ | $1/s$ | Unbinding rate constant | - | $8 \times 10^{-3}$ | 8 |

| | | | | | |
|---|---|---|---|---|---|
| $k_{df}$ | $1/s$ | Degradation rate constant | - | $1\times10^{-6}$ | 12 |
| $k_{int}$ | $1/s$ | Internalization rate constant | - | $5\times10^{-5}$ | 8 |
| $\varphi$ | - | Tumor volume fraction available to drugs | - | $0.4$ | 8 |
| $C_{rec}$ | $M$ | Concentration of cell-surface receptors | - | $1\times10^{-5}$ | 8 |
| $\omega_{Dox}$ | $m^3/mole$ | Survival constant of cancer cells | - | $0.6603$ | 9, 13 |

* This value is only used for validation of fraction of killed cells in classical chemotherapy.

**TABLE SIII.** Parameters used for targeted nano-sized drug delivery system (immunochemotherapy).

| **Parameter** | **Unit** | **Definition** | **Value** | | **Ref.** |
|---|---|---|---|---|---|
| $D_n$ | $m^2/s$ | Diffusion coefficient of nanocarrier | - | $7\times10^{-12}$ | 8 |
| $D_{eff}$ | $m^2/s$ | Diffusion coefficient of free drug | Tumor | $3.4\times10^{-10}$ | 8 |
| | | | Healthy | $1.58\times10^{-10}$ | |
| $K_d$ | $min$ | Blood circulation decay constant | - | $600$ | 8 |
| $k_{on}$ | $1/(M\,.s)$ | Binding rate constant | - | $1.5$ | 8, 10, 11 |
| $k_{off}$ | $1/s$ | Unbinding rate constant | - | $8\times10^{-3}$ | 8 |
| $k_{int}$ | $1/s$ | Internalization rate constant | - | $5\times10^{-5}$ | 8 |
| $k_{rel}$ | $1/s$ | Release rate constant | - | $2.1\times10^{-6}$ | 8 |
| $k_{out}$ | $1/s$ | Free payload efflux rate from cells | - | $5.52\times10^{-5}$ | 14-16 |
| $k_{in}$ | $1/s$ | Free payload uptake rate into cells | - | $2.31\times10^{-3}$ | 14, 15 |
| $\varphi$ | - | Tumor volume fraction available to drugs | - | $0.05-0.1$ | 8 |
| $\varphi^+$ | - | $Ag^+$ cell volume fraction | - | $0.7$ | 14 |

| | | | | | |
|---|---|---|---|---|---|
| $\varphi^{-}$ | - | $Ag^{-}$ cell volume fraction | - | 0.3 | [14] |
| $\alpha$ | - | Number of particles in each nanocarrier | - | 100 | [8] |

**TABLE SIV.** Boundary conditions used in computational modeling of drug delivery systems.

| Regions | Boundary conditions | |
|---|---|---|
| | Interstitial fluid flow | Solute transport |
| Inner boundary | $(-\kappa_t P_i\|_{\Omega^t}) = (-\kappa_n P_i\|_{\Omega^n})$<br>$(P_i\|_{\Omega^t}) = (P_i\|_{\Omega^n})$ | $\left((D_{eff}{}^{t}\nabla C - V_i C)C\big\|_{\Omega^t}\right) = \left((D_{eff}{}^{n}\nabla C - V_i C)C\big\|_{\Omega^n}\right)$<br>$(C\|_{\Omega^t}) = (C\|_{\Omega^n})$ |
| Outer boundary | $P_i = \text{Constant}$ | $-\text{n}.\nabla C = 0$ |

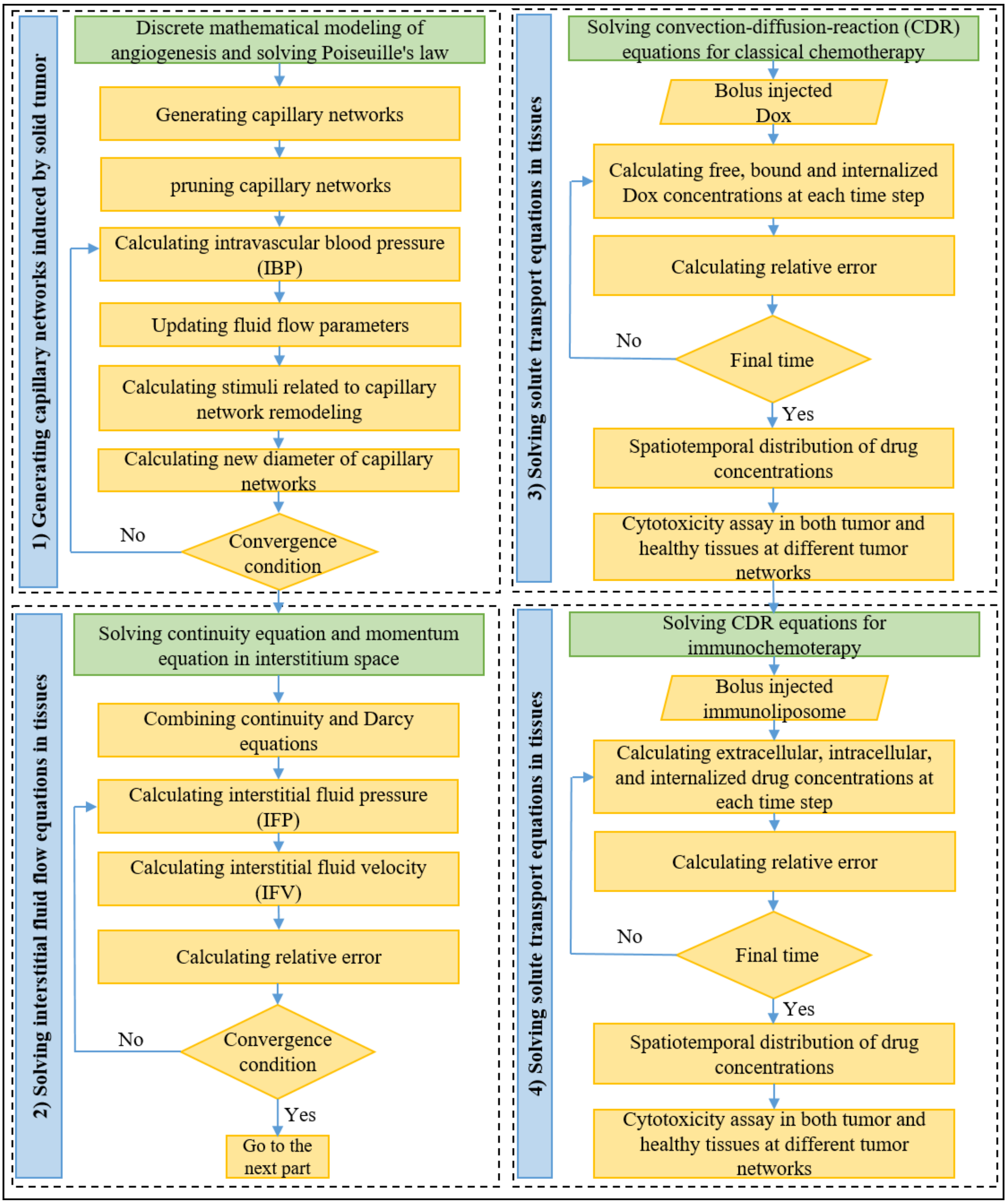

**FIG. S1.** Block diagram illustrating different computational approaches used to build the present comprehensive model. Further details are contained in the solution strategy and computational modeling section of the main text.

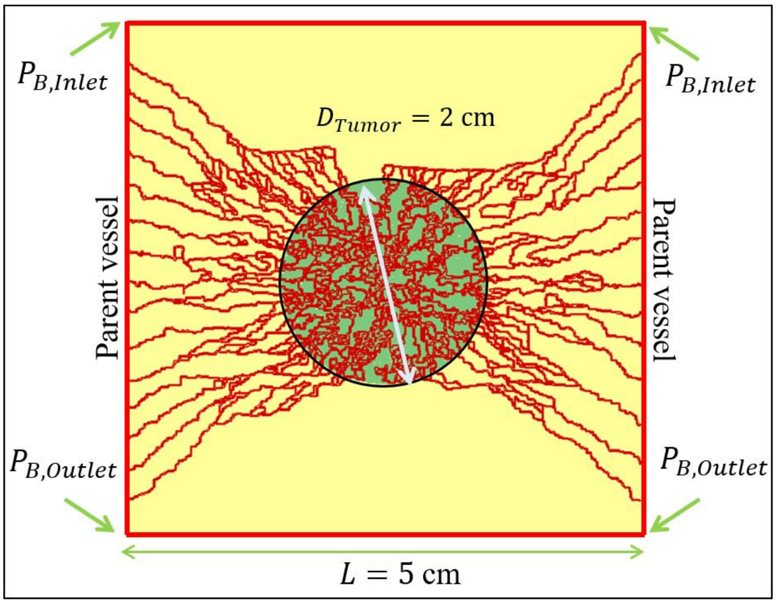


**FIG. S2.** Schematic of computational domain consisted of the solid tumor, surrounding healthy tissue, and microvascular networks. Boundary conditions used in intravascular blood flow simulation are also shown.

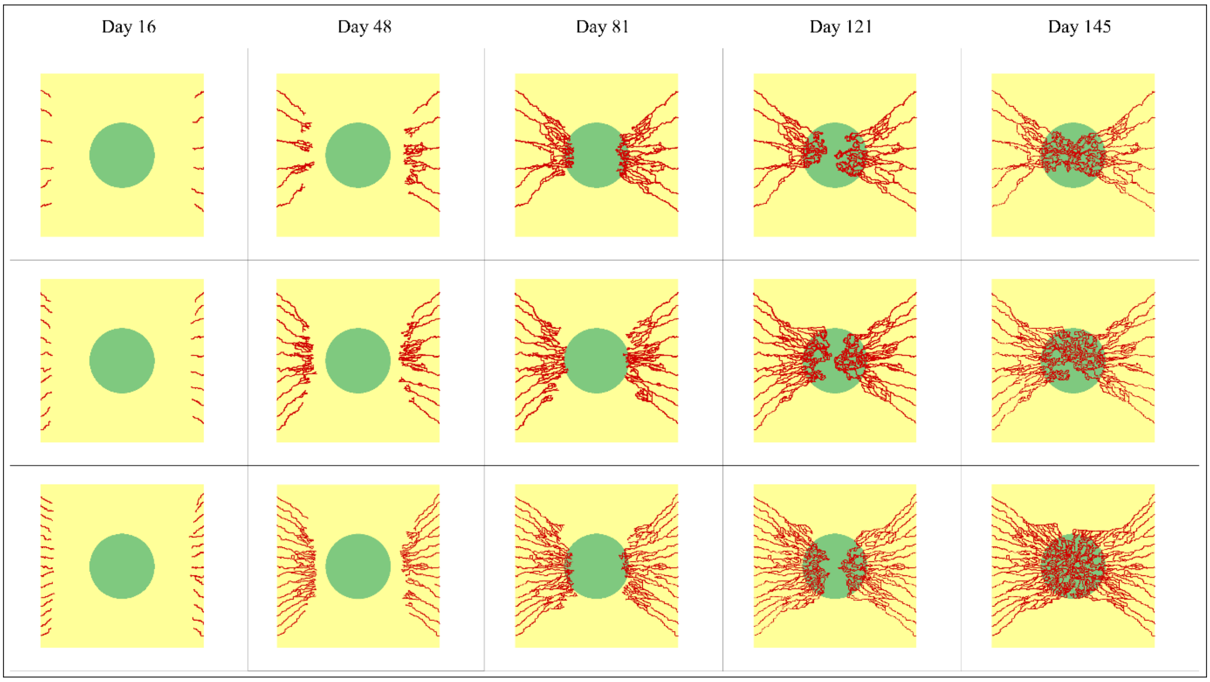


**FIG. S3.** Construction of microvascular networks in a 2-cm tumor at days 16, 48, 81, 121, and 145 of tumor-induced angiogenesis. The microvessels grow from 6-15 initial capillary sprouts on the parent vessels. The solid tumor, which is placed at the center of the computational domain, secretes vascular endothelial growth factor (VEGF) which diffuses into the tissue and then assists in the direction of movement of the endothelial cell into the tumor.
